# A composite filter for low FDR of protein-protein interactions detected by in vivo cross-linking

**DOI:** 10.1101/2020.05.15.097956

**Authors:** Luitzen de Jong, Winfried Roseboom, Gertjan Kramer

## Abstract

In vivo chemical cross-linking combined with LCMSMS of digested extracts (in vivo CX-MS) can reveal stable and dynamic protein-protein interactions at a proteome wide-scale and at the peptide level. In vivo CX-MS requires a membrane permeable and cleavable cross-linker that enables isolation of target peptides and a fast and sensitive search engine to identify the linked peptides. Here we explore the use of the search engine pLink 2 for analysis of a previously obtained LCMSMS dataset from exponentially growing *Bacillus subtilis* treated in culture with the cross-linker bis(succinimidyl)-3-azidomethyl-glutarate (BAMG). Cross-linked peptide pairs were identified by pLink 2 in very short time at an overall FDR of < 5%. To also obtain a FDR < 5% for inter-protein cross-linked peptide pairs additional thresholds values were applied for matched fragment intensity and for the numbers of unambiguous y and b ions to be assigned for both composite peptides. Threshold values were based on a set of decoy sequences from yeast and human sequence databases. Also the mass- and charge-dependent retention times of target peptides purified by diagonal strong cation exchange chromatography were used as a criterion to distinguish true from false positives. After this filtering, pLink 2 identified more than 80% of previously reported protein-protein interactions. In addition the use of pLink 2 revealed interesting new inter-protein cross-linked peptide pairs, among others showing interactions between the global transcriptional repressor AbrB and elongation factor Tu and between the essential protein YlaN of unknown function and the ferric uptake repressor Fur.

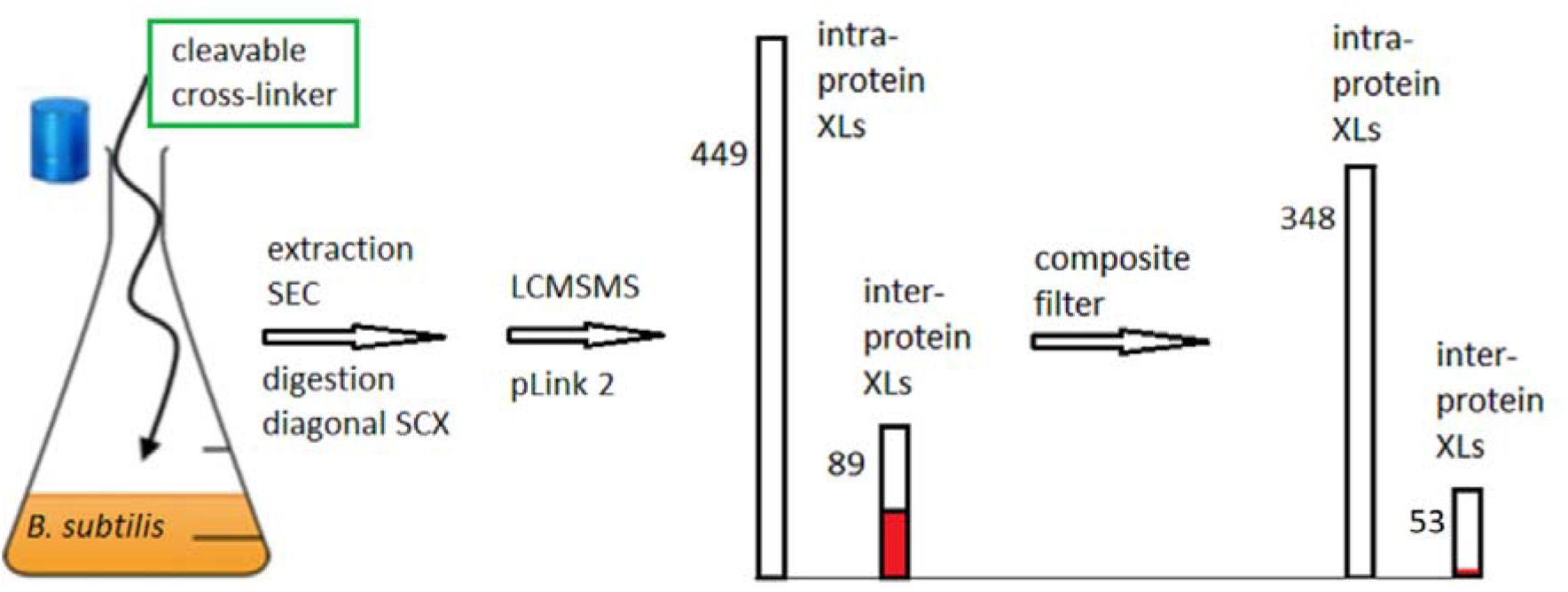

**Highlights:**

- Improved protocol for identification of PPIs at low FDR by in vivo cross-linking with BAMG
- The use of all intra-protein cross-linked peptide pairs as true positives
- The cytosolic aminopeptidase (AMPA_BACSU) interacts with the 50S ribosomal protein L17
- The transition state regulator AbrB interacts with elongation factor Tu
- The essential protein YlaN of unknown function interacts with the iron uptake repressor Fur

**Significance:** Important for reliable identification of PPIs by chemical cross-linking in vivo is a low FDR of non-redundant inter-protein peptide pairs. Here we describe how to recognize the presence of spurious interactions in a dataset of cross-linked peptide pairs enriched by 2D strong cation exchange chromatography and identified by LCMSMS by taking into account chromatographic behavior of cross-linked peptide pairs and protein abundance of corresponding peptides. Based on these criteria we assessed that the FDR of the fraction of non-redundant inter-protein cross-linked peptide pairs was approx. 20-25% by interrogating an entire species specific database at an overall FDR of 5% or 0.1% with a search engine that otherwise scores best in sensitivity among other search engines. We have defined a composite filter to decrease this high FDR of inter-protein cross-linked peptide pairs to only about 2%.

## 1. Introduction

Specific protein-protein interactions (PPIs) are crucial for the regulation of biochemical processes. Large scale approaches like affinity purification combined with mass spectrometry[1], proximity-dependent labeling[2], the use of co-elution profiles[3] and the yeast two-hybrid system[4] can reveal which proteins interact with each other under different experimental and physiological conditions. In vivo chemical cross-linking followed by mass spectrometry and database searching to identify cross-linked peptide pairs (CX-MS) has the potency to reveal cellular protein-protein interactions at a proteome-wide scale in a single experiment and in a short time[5–7]. Besides stable protein complexes also dynamic assemblies that may dissociate upon cell extraction can be trapped by cross-linking in vivo, implying that PPIs may be encountered that have thus far escaped detection by in vitro approaches. Most importantly, the spatial arrangement of proteins in a complex can be assessed by CX-MS by virtue of the identity of linked amino acid residues in the protein sequences as determined by MS and the known length of the spacer of the cross-link. The presence of cross-links formed between two proteins in a complex facilitates modeling of the overall structure if the coordinates of the atoms in the two interacting proteins are known. Such models may lead to hypotheses about the functional significance of the interaction. Thus, CX-MS in vivo is a useful approach in structural and systems biology.

In vivo CX-MS requires a membrane permeable cross-linker designed to facilitate mass spectrometric identification of cross-linked sites. Peptide identification is achieved by liquid chromatography coupled to mass spectrometry (LCMSMS). Mass and charge of the peptides are determined in the MS1 stage. In the MS2 stage peptides are selected for fragmentation by cleavages of the peptide bonds induced by collision with gas molecules (collision-induced dissociation, CID) in a data dependent way, i.e., selection is dependent on signal intensity, mass and charge, and whether a particular precursor ion has been selected before in a given time window. This results in an MS2 fragment spectrum that is characteristic for the peptide sequence. With an MS1MS2 dataset, unmodified peptides, or peptides with a defined modification, can be identified by searching in a sequence database. However, for cross-linked peptides this approach is challenging for two main reasons. In the first place the MS1 data give no information about the masses of the composite peptides, which hampers their identification by database searching. The lack of peptide mass knowledge can be circumvented using a gas phase cleavable cross-linker yielding fragment ions from which the masses of the composite peptides can be deduced. In the second place cross-linked peptides are present in sub-stoichiometric amounts as compared with unmodified peptides, so that selection in the mass spectrometer in the process of data dependent acquisition of MS1MS2 spectra is hampered. This problem requires enrichment of the rare cross-linked species out of the bulk of unmodified peptides.

To meet these requirements and challenges we previously synthesized bis(succinimidyl)-3-azidomethyl-glutarate (BAMG)[8], a membrane permeable cross-linker[9]. The short spacer of BAMG gives relatively high resolution cross-link maps, while the azido group in the spacer can be modified in different ways. This reactive versatility enables three different cross-link analysis strategies[10–14]. For CX-MS in vivo we have made use of a TCEP-induced reduction of the azido group to enrich cross-linked peptide pairs obtained from growing *Bacillus subtilis* cells treated in culture with BAMG[9]. Besides isolation of target peptides, reduction of the azido group also facilitates mass spectrometric identification of the linked peptide pair. This is due to the fact that the two cross-link amide bonds of a peptide pair are cleavable in the gas phase by CID [11]. The cleavage of the cross-link amide bond can occur along with a peptide bond cleavage. The principle was demonstrated with protein complexes from *Bacillus subtilis* in the mass range 400 kDa to 1-2 MDa obtained by size exclusion chromatography (SEC) after in vivo cross-linking. We identified several cross-linked peptides revealing transient and stable PPIs of high biological significance and low FDR by searching MS1MS2 data from the entire *B. subtilis* sequence database[9]. This analysis was supported by the use of in-house developed scripts called Raeng and YeunYan in combination with the search engine MASCOT.

This procedure, further called the Raeng/Mascot approach, is relatively laborious, making it unattractive for general use. This holds in particular in cases were large datasets are to be expected by the use of state of the art HPLC coupled to sensitive and fast mass spectrometers like the recently introduced trapped ion mobility time-of flight mass spectrometer (timsTOF Pro) enabling online parallel accumulation-serial fragmentation (PASEF)[15]. Here we explore the use of the recently launched second generation search engine, pLink 2[16]. In comparison with many other search engines developed for cross-link analysis, pLink performs best with respect to number of identified cross-links with good FDR at cross-link spectrum matches (CSM) level [16,17]. Moreover it is extremely fast, and is also suitable for analysis using cleavable cross-linkers like BAMG[16]. With an existing LCMSMS dataset we observed a large overlap with the results obtained with the Raeng/Mascot approach, but also found evidence that the FDR for inter-protein peptide pairs is higher for inter-protein peptide pairs than for intra-protein peptide pairs. Here we describe an approach to further lower the FDR for the latter type of cross-links that reliably led to interesting new protein-protein interactions.

## 2. Materials and methods

### 2.1. Acquisition of the LCMSMS dataset to explore the use of pLink 2

The LCMSMS dataset was acquired as described previously in detail [9]. In short, exponentially growing *B. subtilis* (strain 168) was treated in culture with 2 mM BAMG at 37° C for 5 min. The soluble extract, obtained after harvest and sonication of cross-linker-treated cells, was subjected to SEC. A tryptic digest of protein complexes in the size rage 400 kDa to 1-2 MDa was selected for further analysis. First the peptide mixture was subjected to diagonal strong cation exchange (SCX) chromatography to sequester the cross-linked peptides from the bulk of unmodified peptides. Fractions enriched in cross-linked peptide pairs were analyzed by LCMSMS using an Eksigent Expert nanoLC 425 system connected to the Nano spray source of a TripleTOF 5600+ mass spectrometer. Data processing was as described [9]. For analysis by pLink 2 the original mgf files were combined into 13 mgf files. An overview of the origin and size of the combined mgf files along with the mass accuracy used for pLink 2 searches is shown in Supplementary Table S1. The original mgf files are available via ProteomeXchange with identifier PXD006287.

### 2.2. Identification of cross-linked peptides

BAMG-cross-linked peptides become cleavable in the gas phase when the azido group in the spacer of the cross-link has been reduced to an amino group. For cross-link identification we used pLink 2 as a search engine operating in the stepped-HCD mode for MS-cleavable cross-linkers [16]. This mode has been developed for the cross-linkers DSBU[18] and DSSO[19]. In the stepped-HCD mode cleavage of the DSSO- or DSBU-cross-link and generation of peptide fragments occurs with two different CID energies after which the data obtained in the two steps are combined in one file entry. Cleavage of the BAMG-cross-link amide bonds and cleavage of the peptide bonds resulting in y and b ions occurs in one step under defined CID conditions (Fig 1). Therefore both DSSO-, DSBU- and BAMG-cross-linked α-β peptide pairs yield similar MSMS spectra containing the signals from intact cleaved peptides along with the signals from peptide bond cleavages. Like DSSO and DSBU, BAMG is scissile at two identical sites. This implies that a single cleavage event in the members of an ensemble of identical cross-linked peptide pairs results in a mixture of two pairs of products with a characteristic mass difference. The short versions of the cleaved peptides are denoted αS and βS and the long version are denoted αL and βL. For BAMG the mass difference between αS and αL and between βS and βL is 125.048 Da, αL and βL being modified by the remnant of the cross-linker in the form of a γ-lactam, while αS and βS are unmodified after the cleavage (Fig 1). The presence of four such cleavage products directly reveals the masses of the composite peptides. Also double cleavage events usually take place, by which cleavage of a cross-link amide bond occurs along with a peptide bond cleavage. Often signals from only one pair of cleavage products are detectable, usually from the shortest composite peptide, the other pair of cleavage products being completely fragmented by secondary cleavages. If only one pair of peaks with a 125 Da mass difference is present in the mass spectrum, the mass of the other peptide can be calculated by subtracting the mass of the first peptide from the mass of the precursor ion. We have shown that 83% of the mass spectra from 401 different cross-linked peptide pairs display at least one pair of cleavage products from either the α-peptide or the β-peptide. In principle the presence of 2 cleavage products in the combinations αS and βS, αS and βL, αL and βS and αL and βL can also be used to deduce the masses of α and β. These combinations occur in an additional 5% of the instances [11]. Peptide bond cleavages can also result in ion pairs differing 125.048 Da, denoted αSy or αSb for the short and αLy or αLb for the long (+ 125.048 Da) versions of y or b ions from the α peptide. For the β peptide short and long version of y and b ions are denoted βSy, βLy, βSb and βLb. pLink 2 was adapted for BAMG cross-link identification by considering BAMG-specific fragment ions. It is anticipated that the presence of y-ion or b-ion pairs with mass differences of 125.024 Da will not prevent assessment of the correct masses of α and β by pLink 2, although the presence of these ion pairs may result in the calculation of more than one candidate for the masses of α and β of a given precursor ion.

**Figure 1.**
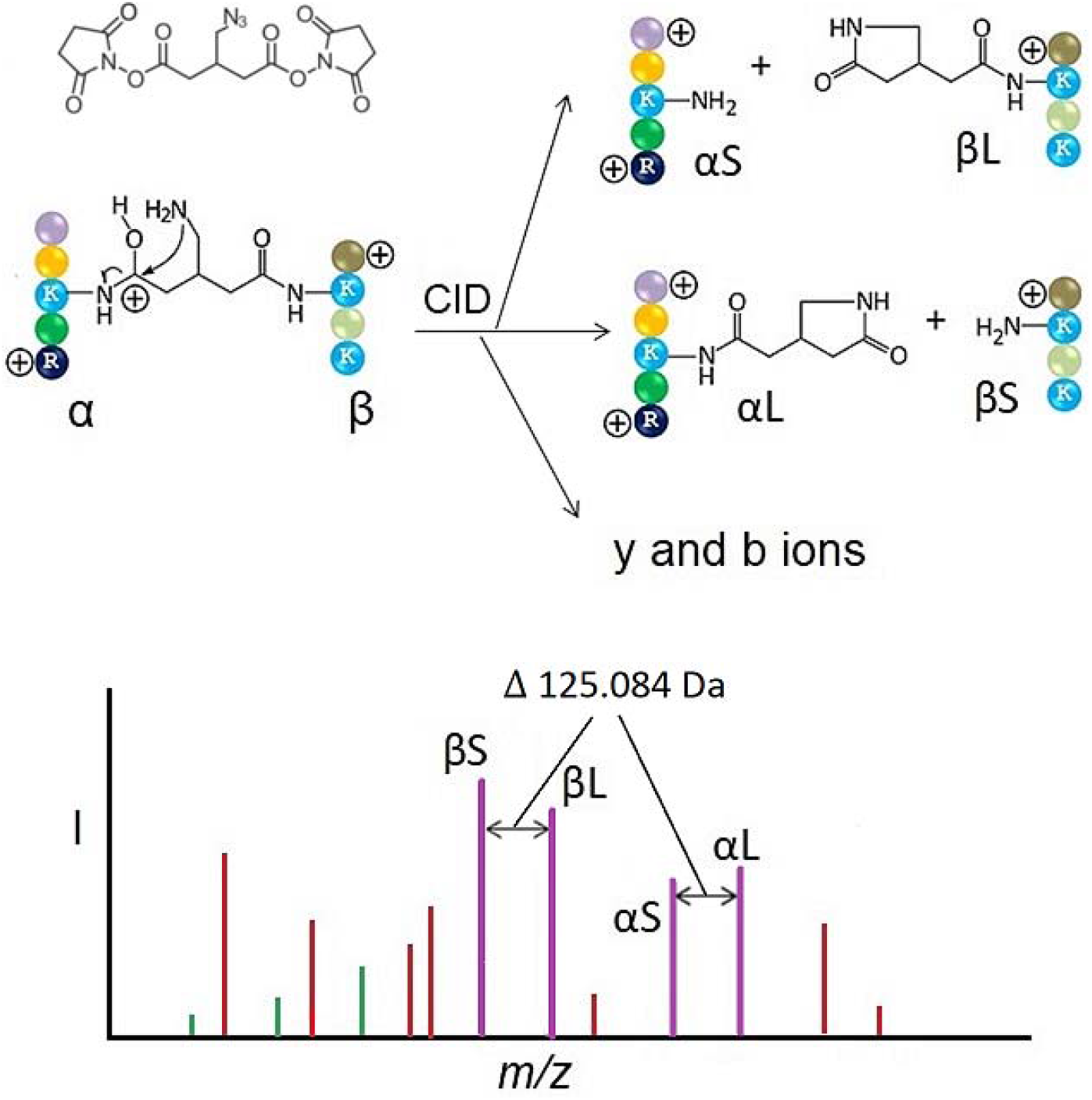
Gas phase cleavage reactions of BAMG-cross-linked peptides in which the azido group has been reduced to an amine group. Upper left corner, structure of BAMG. Middle part, collision induced dissociation (CID) of a cross-linked peptide pair leads to cleavages of the two cross-link amide bonds along with cleavages of peptide bonds resulting in y an b ions. Cleavage of an amide bond probably occurs by nucleophilic attack of the amine in the spacer of the cross-link to a protonated carbonyl group of the amide bond. This leads to formation of an unmodified peptide or short version of the cleavage product (αS or βS), the other peptide being modified by the remnant of the cross-linker in the form of a γ lactam adding 125.048 Da to the mass of the peptide. This is the longer version of the cleavage product (αL or βL). Amino acids are depicted as colored candies. The indicated gas phase charge states of the cross-linked peptide and the cleavage products are arbitrarily. The lower part is a cartoon of a fragment mass spectrum with two pairs of cleavage products with the characteristic 125.048 Da mass difference (purple sticks) and some peaks of b (green) and y (red) ions.

Four different protein databases were used for identification; (i) the Uniprot protein database of *Bacillus subtilis* (4260 entries); (ii) a hybrid database composed of the proteins from database (i) and the Uniprot protein database of Saccharomyces cerevisiae (6043 entries) [20]; (iii) a database of the 673 proteins identified in the primary SCX fractions and (iv) a hybrid database composed of the proteins from database (iii) and 1085 human proteins identified in a SEC fraction of a HeLa cell nuclear extract[12]. The yeast and human proteins in the hybrid databases are used as a source of decoy sequences for FDR estimations.

### 2.3. Mass spectrometric criteria for cross-link identity assignment

The following parameters were used for pLink 2 searches of tryptic peptides using the mgf dataset as shown in Supplementary Table S1; mass accuracy for precursor and fragment ions: 25-75 ppm, depending on the mgf file; up to two missed cleavages allowed; masses: 600-6000; lengths for α and β peptides: 6-60 amino acids; a carbamidomethyl group at C as fixed modification; oxidation at M as variable modification; 5% or 0.1% FDR for cross-linked spectrum matches. With these settings intra-protein cross-links were assigned as nominated by pLink 2. For assignment of inter-protein cross-links the following additional requirements were taken into account: the assignment of least 4 unambiguous y ions for α and β peptides with a length of 12 amino acids or more, and at least 3 unambiguous y ions for peptides consisting of 11 amino acids or less along with a matched intensity scoring higher than 35%. For a matched intensity score of more than 50%, at least 2 unambiguous y ions are sufficient for assignment of peptides consisting of 11 amino acids or less, provided that also at least 2 unambiguous b ions can be assigned to the peptide. A y or b ion is considered ambiguous if it can also be assigned to one or more other fragments. A yS and yL or bS and bL ion pair with the 125 Da mass difference is counted only once for the requirement with respect to the minimal number of unambiguous y or b ions for validation and assignment. Doubly charged y or b ions at m/z ≤ 700 are not taken into account. The ignorance of b ions as a selection criterion, except for short peptides and at high matched intensity as described above, is based on their relatively low occurrence as compared with y ions [9].

Matched intensity and numbers of unambiguously assigned y and b ions are mentioned in result tables only once for candidates of which more than one MSMS result was put forward as a result of multiple selections for MSMS by the mass spectrometer. Usually the MSMS spectrum with the highest matched intensity was chosen, provided that also the other criteria for assignment were met.

### 2.3. Elution time window during SCX as a criterion for cross-link identity assignment

A further filter concerned the mass of a cross-linked peptide pair and the calculated charge state in relation to the elution time. The charge state is calculated at pH 3, i.e., the conditions of strong cation exchange chromatography, assuming protonation of all acid and basic groups. To asses the distribution of mass and charge in relation of elution time in SCX chromatography of true positives we took into account all intra-protein cross-linked peptide pairs identified by pLink 2 and only the inter-protein peptide pairs that were identified by both Raeng and pLink 2. Identical peptide pairs that eluted in different SCX fractions were also taken into account.

### 2.4. FDR estimation; the use of different decoy sequences for inter- and intra-protein peptide pairs; selectivity and sensitivity

The overall FDR is defined as FDR = d/(d + t) × 100%, in which d is the number of decoy α-β sequence hits and t is the number of identified target α-β sequences. A decoy α-β peptide pair consists of two human or yeast sequences or of one human or yeast and one target (*B. subtilis*) sequence. For intra-protein cross-linked peptide pairs, decoy sequences for α and β are from the same yeast or human protein, whereas for inter-protein peptide pairs the decoy sequences are from different proteins. Also target-reversed and reversed-reversed peptide pairs with different sequences of α and β peptides from the same proteins are decoy sequences for intra-protein cross-linked peptide pairs. However, previously we showed that no reversed-reversed versions and only three target-reversed versions of these intra-protein decoy sequences were put forward on a total of 1288 decoy hits[11].

## 3. Results and discussion

### 3.1. Protein composition of the extract obtained after in vivo cross-linking of exponentially growing cells with BAMG

Previously we have identified cross-linked peptide pairs from protein complexes in the size range 400 kDa to 1-2 MDa after SEC of an extract obtained from an exponentially growing *B. subtilis* culture treated with BAMG. A size exclusion chromatogram of the protein extract is shown in Fig 2.

**Figure 2.**
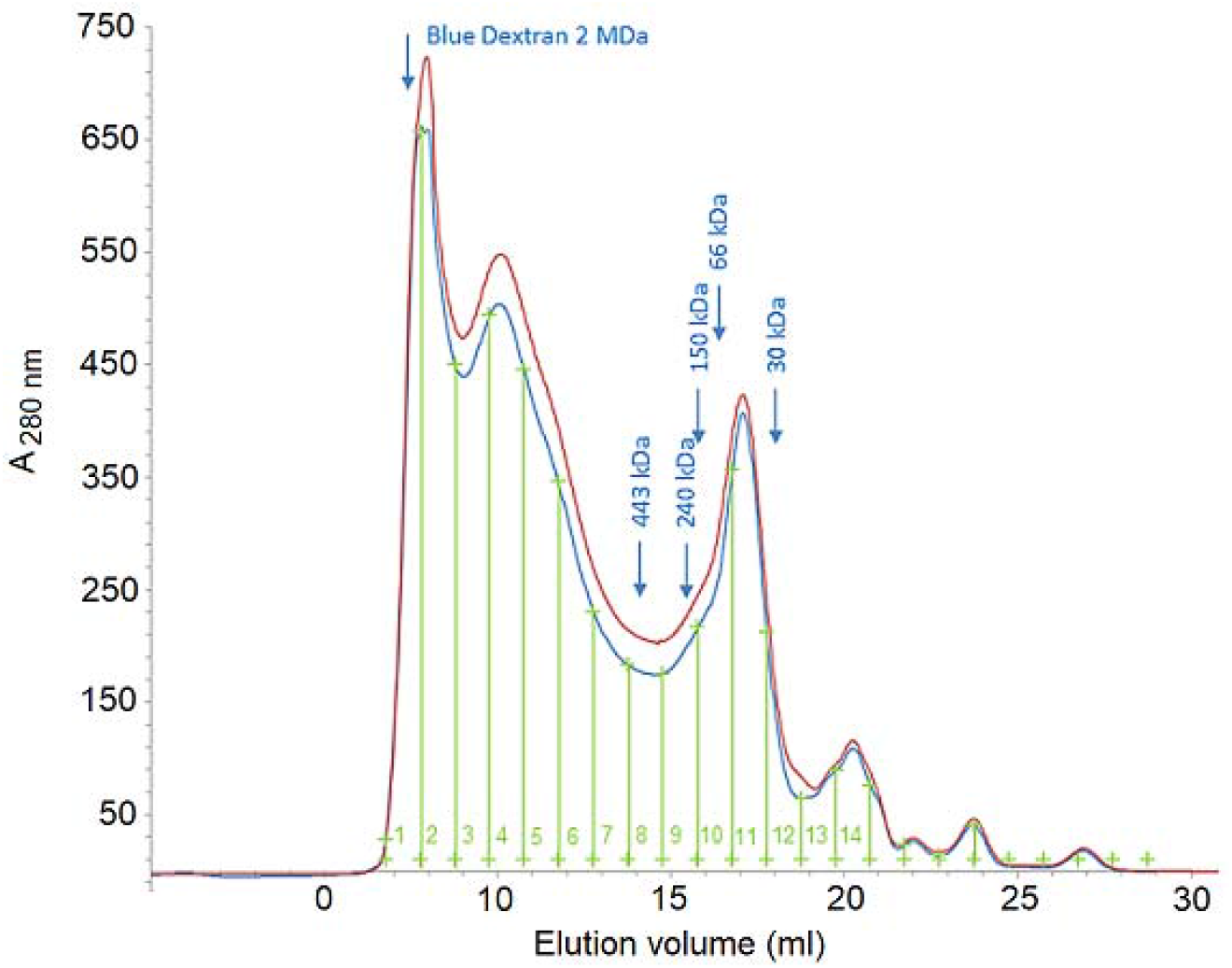
Size exclusion chromatogram on a Superose 6 10/300 GL column of a soluble extract from Bacillus subtilis cross-linked in vivo with BAMG. MW markers, 443 kDa, apoferritine (horse spleen); 240 kDa catalase (bovine liver); 150 kDa, alcohol dehydrogenase (yeast); 66 kDa, bovine serum albymine, 30 kDa carbonic anhydrase (bovine erythrocytes). Fractions are indicated by green vertical lines. Duplicate chromatograms are shown by red and blue lines. Material in fraction 4-7 was used in this study.

In the selected SEC fractions 4-7 we identified 673 proteins [9]. The known MW of most of these proteins is much smaller that the size range of 400 kD to 1-2 MDa. Besides specific cross-linking in stable and transient protein complexes also cross-linking during random diffusional encounters between proteins in the concentrated cytosol may attribute to this high molecular weight shift and size heterogeneity. It is anticipated that only specific interactions will be found, assuming that in a digest the many particular non-specific cross-links are so rare that they will remain largely undetected by LCMSMS.

### 3.2. Isolation of cross-linked peptides by diagonal strong cation exchange (SCX) chromatography

The principle of diagonal chromatography to isolate BAMG-cross-linked peptides from the bulk of unmodified peptides [12] is depicted in Fig 3. Isolation of peptides with a specific reactivity by two-dimensional liquid chromatography was introduced in 2002 in order to increase the numbers of proteins identified by LCMSMS [21]. In this approach, coined diagonal chromatography, target peptides in chromatographic fractions are subjected to a reaction that modifies their chromatographic retention time. In the second dimension peptides in treated fractions sequester from the bulk of unmodified peptides that elute at the same retention time as in the primary run. Here we use reduction of the azido group to an amino group on the spacer of a crosslink in peptides to increase the charge state. Also peptides with an internal cross-link, called loop-linked peptides and peptides with one modified lysine residue, called mono-linked peptides, the second reactive ester of the cross-linker being hydrolyzed, are enriched by diagonal SCX chromatography.

**Figure 3.**
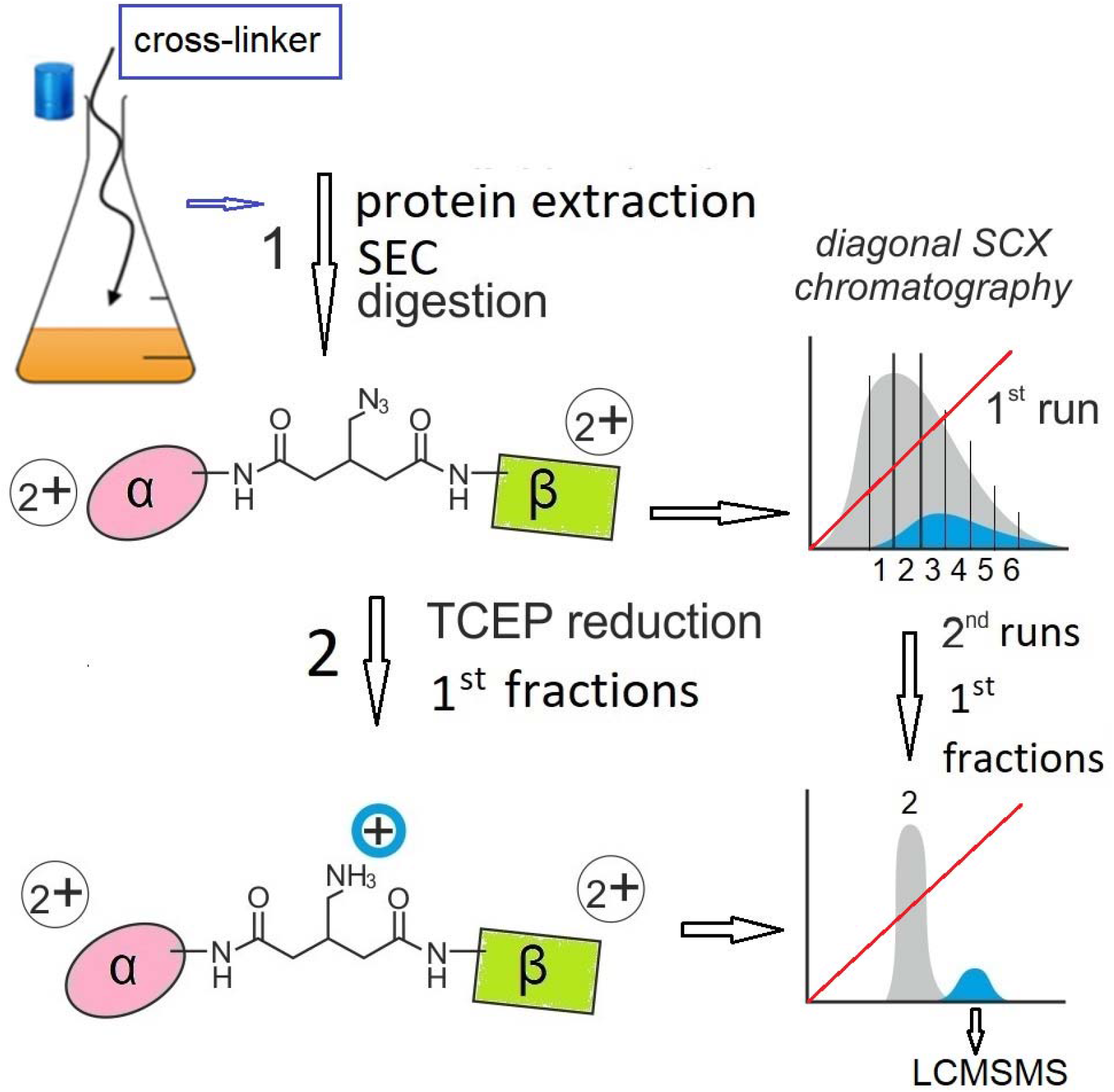
Workflow from in vivo cross-linking with BAMG to LCMSMS. Left part, 5 min after addition of BAMG to an exponentially growing Bacillus subtilis culture the cross-linker is quenched and cells are harvested and sonicated. (1), the cross-linked protein extract is subjected to SEC and then digested to obtain a peptide mixture with cross-linked α-β peptide pairs. Right part, the peptide mixture is fractionated by strong cation exchange chromatography (first dimension SCX), using a mobile phase of pH 3 and a salt gradient (red lines) of ammonium formate to elute bound peptides. Grey, regular peptides; cyan, cross-linked peptides. (2), reduction by TCEP of the azido group in the spacer of the cross-linker to an amine group in selected SCX fractions. At the pH of the mobile phase of SCX chromatography the amino group is protonated adding an extra positive charge to cross-linked peptides. The TCEP-treated primary fractions are then separately subjected to the secondary runs of SCX. Here target peptides are sequestered from the bulk of unmodified peptides that elute at the same time as in the primary run, while the extra charge state of the cross-link peptides leads to elution at a later time. Depicted peptide charge states after (1) and (2) are calculated for pH 3, assuming full protonation of the two amino-termini plus 2 basic amino-acid side chains, carboxylic acid side chains being uncharged under these conditions.

An additional advantage of purification of target peptides by diagonal SCX chromatography is the characteristic relationship between de elution time and the mass range and positive charge range of cross-linked peptide pairs at pH 3.0 of the mobile phase (Table 1). Table 1 has been composed from the data of 626 cross-linked peptide pairs (section 2.4), showing that (i) highly charged cross-linked peptide pairs tend to elute late and (ii) at each charge state 4^+^, 5^+^ and 6^+^ the mass range decreases at increasing elution time. Three 5^+^ charged peptides from fraction 10 and one 6^+^ charge peptide from fraction 12 form the only exceptions to the otherwise consistent distribution. This consistency enables the use of elution time in relation to the charge and mass of a candidate cross-link as one of the criteria to discriminate between true and false positives.

**Table 1.** Charge at pH 3 and mass distributions of cross-linked peptide pairs in SCX fractions

| SCX<br>frac-<br>tion | XL<br>pep-<br>tide<br>pairs<br>(n) | Mass range<br>(Da) at<br>charge +3 | XL<br>pep-<br>tide<br>pairs<br>(n) | Mass range<br>(Da) at<br>charge +4 | XL<br>pep-<br>tide<br>pairs<br>(n) | Mass range<br>(Da) at<br>charge +5 | XL<br>pep-<br>tide<br>pairs<br>(n) | Mass range<br>(Da) at<br>charge 6+ | XL<br>pep-<br>tide<br>pairs<br>(n) |
| --- | --- | --- | --- | --- | --- | --- | --- | --- | --- |
| 7 | 30 | 1793-1868 | 2 | 2274-4505 | 28 |  |  |  |  |
| 8 | 62 | 1869-2109 | 2 | 2274-4387 | 59 | 4377 | 1 |  |  |
| 9 | 212 |  |  | 1756-3890 | 201 | 3273-4672 | 11 |  |  |
| 10 | 72 |  |  | 1641-3033 | 69 | 2554-3394 | 3 |  |  |
| 11 | 83 |  |  | 1734-2858 | 45 | 2861-5181 | 36 | 4055-6092 | 2 |
| 12 | 57 |  |  | 1513-2568 | 17 | 2540-4462 | 39 | 2661 | 1 |
| 13 | 52 |  |  | 1562-2090 | 5 | 2262-3828 | 46 | 3884 | 1 |
| 14 | 39 |  |  | 1435-1779 | 4 | 1967-3351 | 24 | 3419-4919 | 11 |
| 15 | 2 |  |  |  |  | 1862-2595 | 2 | 3405 | 1 |
| 16 | 17 |  |  |  |  | 1862-3510 | 13 | 3313-3870 | 4 |

### 3.3. pLink 2 is a fast search engine for identification of BAMG-cross-linked peptides, but for a FDR < 5% for inter-protein cross-linked peptide pairs further filtering is required

We tested the suitability of pLink 2 as a search engine to identify BAMG-cross-linked peptides using an existing LCMSMS dataset. The dataset consisted of mgf files from LCMSMS analysis of 10 fractions obtained by diagonal SCX chromatography of trypsin-digested material present in SEC fraction 4-7. Previously we identified with the Raeng/Mascot approach several inter-protein cross-linked peptide pairs that fulfill the requirements of a minimal peptide length of 6 amino acids for identification by pLink 2 (Table 2). These inter-protein cross-linked peptide pairs revealed 41 interactions between different proteins. The dataset also contained several homo-dimeric peptide pairs and intra-protein peptide pairs. Although the results were obtained by searching the entire *B. subtilis* sequence database, all inter-protein cross-linked peptide pairs belong to the set of 673 actually identified proteins in the analyzed SEC material. Only a few homo-dimeric and intra-protein peptide pairs did not belong to the 673 identified proteins.

**Table 2.**
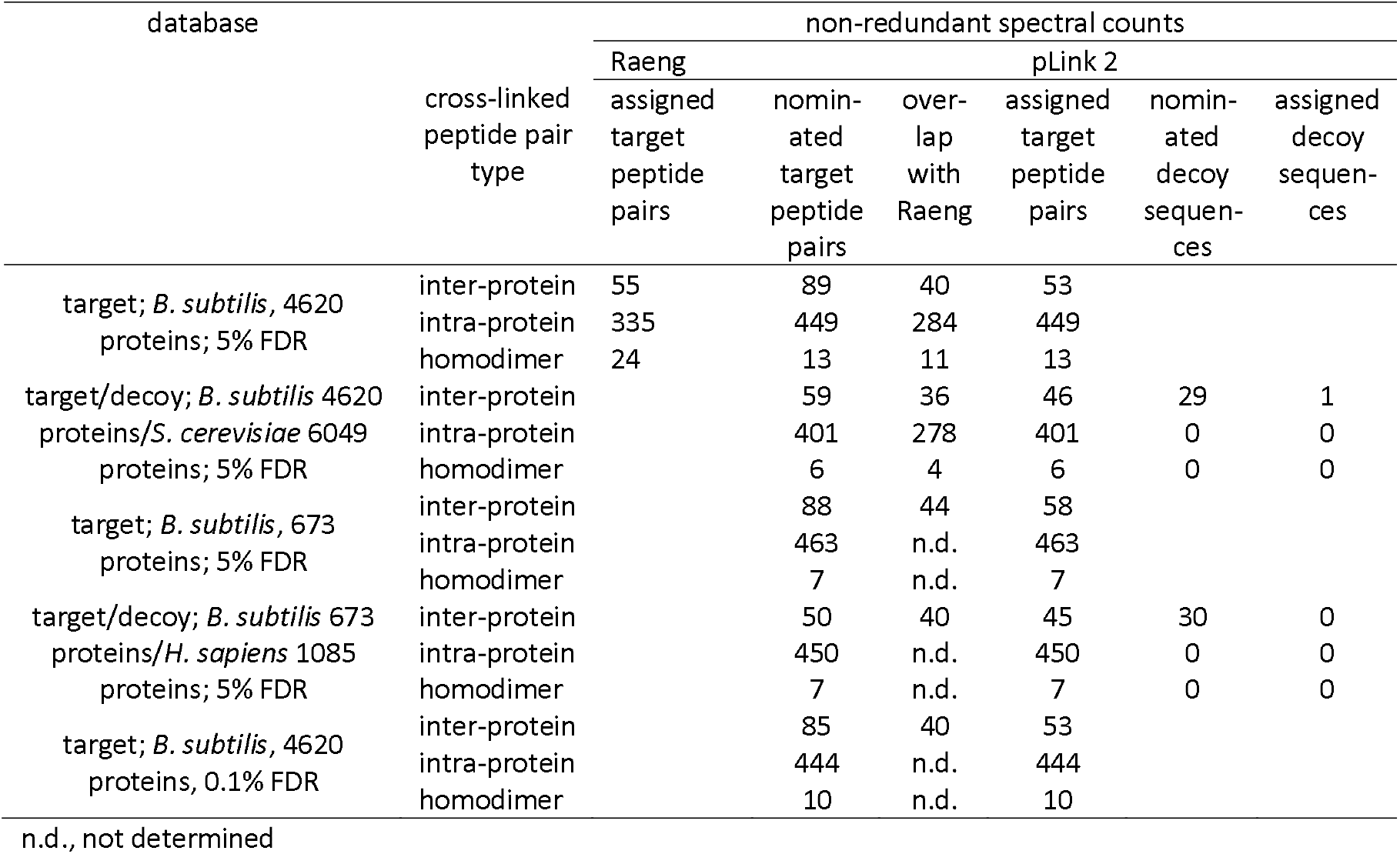
Overview of non-redundant spectral counts of nominated and assigned cross-linked peptide pairs

The 13 MSMS data files (supplementary Table S1) in mgf format with an average size of about 14.4 MB were separately analyzed by pLink in the very short time of about 3 min on average per mgf file. Nearly 10,000 spectra were identified by pLink 2 at less than 5% FDR (Table 3), distributed over intra-protein and inter-protein cross-linked peptide pairs, mono-linked peptides, loop-linked peptides and regular peptides. Table 2 depicts the numbers of the non-redundant intra-protein cross-linked peptide pairs, inter-protein peptide pairs and homo-dimeric peptide-pairs identified by pLink 2. Less than 3% of the intra-protein cross-linked peptide pairs did not belong to the 673 independently identified proteins (Supplementary Table S2, column F, salmon-pink highlighted). However, no less than 24 out of the 89 inter-protein cross-linked peptide pairs nominated by plink 2 did not belong to the 673 identified proteins (Supplementary Table S2, column F, pink and cyan highlighted). Moreover, 16 of the 89 nominated species violated the expected elution time in CX chromatography based on their mass and calculated charge at the pH (pH 3.0) of the mobile phase in SCX chromatography (Table 1 and Supplementary Table S2, column D, blue highlighted). The presence of a relatively large number of inter-protein peptide pair candidates not belonging to the 673 most abundant proteins and the violation by several inter-protein peptide pairs of the mass and charge rules of the SCX elution times strongly suggests the presence of false positive nominations among these inter-protein cross-linked species. This observation is not unexpected, since it is well documented that by searching a large sequence database at a given FDR at CSM level, practically all false positives are confined to inter-protein peptide pairs [9–11]. Based on a total of 1856 spectra of cross-linked peptide pairs (Table 3) we roughly calculate a FDR at CSM level of 1.7% assuming that the 32 total inter-peptide cross-linked peptide pairs with aberrant SCX elution time, or of which at least one peptide does not belong to the 673 independently proteins, are false positives. This would imply a FDR of 11.3% for the 252 inter-protein peptide pairs or, with 30 non-redundant decoy hits, about 25% for the 89 non-redundant inter-protein peptide pairs put forward by pLink 2 (Tabel 2). The FDR would be even larger if related to PPIs. These trends are in agreement with a previous discussion on FDR estimations on different levels[22].

**Table 3.** Overview of spectral counts identified by pLink 2 at <5% FDR

| SCX fraction | Counts |  |  |  |  | Total spectra |
| --- | --- | --- | --- | --- | --- | --- |
|  | Cross-linked spectra |  | Loop-linked spectra | Mono-link spectra | Regular peptide spectra |  |
|  | Intra-protein | Inter-protein |  |  |  |  |
| 7 | 94 | 3 | 183 | 566 | 1185 | 2031 |
| 8 | 124 | 16 | 143 | 478 | 779 | 1540 |
| 9.2 | 12 | 9 | 2 | 30 | 138 | 191 |
| 9.3 | 268 | 39 | 50 | 251 | 161 | 769 |
| 9.4 | 254 | 19 | 103 | 305 | 185 | 867 |
| 9.5 | 32 | 2 | 74 | 136 | 105 | 349 |
| 10 | 174 | 23 | 53 | 150 | 404 | 803 |
| 11 | 229 | 10 | 33 | 172 | 441 | 885 |
| 12 | 173 | 17 | 18 | 130 | 268 | 606 |
| 13 | 146 | 99 | 64 | 133 | 392 | 834 |
| 14 | 66 | 7 | 37 | 47 | 320 | 477 |
| 15 | 6 | 1 | 0 | 4 | 172 | 183 |
| 16 | 26 | 7 | 10 | 22 | 257 | 322 |
| total | 1604 | 252 | 770 | 2424 | 4807 | 9857 |
| percent | 16.3 | 2.6 | 7.8 | 24.6 | 48.8 |  |

### 3.4. A target-decoy database to find criteria for a low FDR of inter-protein peptide pairs

A target-decoy database is used by pLink 2 for FDR calculation. The decoy database consists of the reversed sequence of the target database. Since reversed sequences are not reported by pLink we decided to use a hybrid database in which the entire *B. subtilis* Uniprot protein database (4260 entries) was combined with the entire *Saccharomyces cerevisiae* Uniprot protein database (6043 entries). In this way we could get insight in the nature of false positives, i.e. nominations with one or two yeast sequences, in order to find criteria to discriminate between true and false positives.

The result of a pLink search with the hybrid database is shown in Table 2. With the increase of the search space while keeping the pLink 2 settings at 5% FDR, the numbers of inter-protein and intra-protein peptide pairs overlapping with the Raeng cross-links dropped by about 10% (Table 2). Also the number of homodimeric peptide pairs decreased. In addition, pLink 2 nominated several decoy sequences. All decoy sequences contain peptide pair sequences from different proteins (supplementary Table S3, column F, salmon-pink highlighted). Since no decoy sequences are present with peptide pairs from the same yeast protein, the FDR for intra-protein target peptide pairs must be very low. Previously we also noticed that only about 0.3% of all decoy hits consisted of intra-protein target and reversed sequences in a complex sample of cross-linked human proteins [11]. This justifies the inclusion of all intra-protein cross-links, along with the common set of inter-protein cross-links put forward by Rang and pLink 2, in the dataset to calculate the mass and charge distribution patterns in relation with the elution time in SCX chromatography (Table 1). It appeared that some decoy sequences put forward by pLink 2 eluted in an unexpected time window. We use an anomalous elution time during SCX chromatography as one of the criteria for filtering the dataset to decrease the FDR for inter-protein cross-linked peptide pairs.

We also noticed that many decoy sequences showed relatively few y ions assigned in one or both composite peptides. Furthermore ambiguity in the assignment of y or b ions occurred sometimes. A relatively low matched intensity of MS/MS spectra was also not uncommon. Therefore we applied thresholds for matched intensity and for the required number of unambiguously assigned y ions and sometimes b ions as described in sections 2.3 and 2.4.

Another source of nominated decoy sequences are cases in which pLink 2 put forward 2 candidates for the same precursor ion or for two different precursor ions with the same mass and eluting in the LCMSMS run within a time window of 5 sec. In a few other instances alternative candidates were also possible for a given precursor ion when certain post-translational modifications were taken into account, namely formylation in two instances and carbamidomethylation at the N-terminus in three instances. In supplementary Table S4 these double nominations are listed. In all cases only the best scoring candidate was assigned, or, in case of equal scores, none of the candidates. In supplementary Tables S2, S3, S5 and S6 these co-called ambiguous nominations that are not assigned are cyan highlighted in column C. In future experiments, the otherwise rare carbamidomethylation at the N-terminus can be circumvented by preventing the presence of iodoacetamide during digestion by trypsin. Formylation is probably unavoidable [23] in the sample preparation work flow where relatively high concentrations of ammonium formate are used to elute peptides during SCX chromatography. To keep the formylation level as low as possible, the SCX fractions were snap frozen in liquid nitrogen immediately after elution followed by lyophylisation

The aim of retention time window determination, application of thresholds for matched intensity and for the numbers of unambiguously assigned y and b ions a, and exclusion of double nominations for the same precursor ions is to lower substantially the FDR, while a high percentage of true positive cross-linked pairs should survive the stringent criteria. Since intra-protein cross-links were identified under conditions that no intra-protein decoy sequences were detected, these target cross-links can be considered as true positives. Therefore intra-protein peptide pairs can be used to assess the effect of filtering on the sensitivity of assignment of true positive inter-protein cross-linked peptide pairs.

### 3.5. Application of mass spectrometric and SCX chromatographic criteria to obtain a low FDR for inter-peptide protein pairs

The effect of application of the criteria as formulated in the Material and Methods sections 2.3 and 2.4 and discussed in section 3.4 is shown in Table 2 (column assigned target peptide pairs and column assigned decoy sequences). For details see columns H-J in supplementary Tables S2 and S3). Only one decoy sequence remains that fulfils the criteria, corresponding to a FDR of about 2% for non-redundant inter-protein peptide pairs (Supplementary Table S3). With one exception the nominated inter-protein peptide pairs of which at least one of the α or β peptide did not belong to the 673 independently identified proteins in the sample were all rejected by the filtering (supplementary Table S2). This shows that the composite filter that we developed is very effective to prevent suspicious candidates from assignment.

Application of the composite filter to the intra-protein cross-linked peptide pairs identified by pLink 2 from the entire *B. subtilis* database shows that 77% fulfilled the criteria for assignment of inter-protein peptide pairs (supplementary Table S2, columns G, H and I). This shows that also true positive cross-linked peptide pairs revealing protein-protein interactions can be detected at a very low FDR and high sensitivity.

The small drop in the number of identified peptides when the size of the database increase prompted to interrogate a much smaller database with pLink 2, in the expectation to identify some more cross-linked peptide pairs. In this case the protein database to be searched was composed of the 673 independently identified proteins in the sample. For decoy sequences 1085 human proteins were added to the *B. subtilis* set of proteins. Nominated and identified cross-links and decoy sequences are listed in Supplementary Tables S5 and S6. The results are summarized in Table 2, showing a small increase in the number of assigned inter-protein and intra-protein cross-linked peptide pairs as compared with the results obtained by interrogation of the entire *B. subtilis* sequence database.

### 3.6. Resemblances and differences of results and approaches between pLink 2 and Raeng/Mascot

The combined searches showed that the overlap between the identifications by Raeng/Mascot and pLink 2 is about 80%. In supplementary figure 1 pLabel-generated mass spectra are depicted with input of the mgf files of the inter-protein cross-linked peptide pairs that had escaped detection by pLink 2. All spectra fulfilled the criteria for assignment. On the other hand pLink identified several cross-linked peptide pairs that had been overlooked by Raeng/Mascot (see for 4 examples the mass spectra in supplementary figure 2). The amount of cross-links put forward by pLink 2 and Raeng may depend on how exhaustive pLink 2 searches for the right peptide masses of α and β, sometimes against a background of secondary fragment pairs differing 125.048 Da, and how exhaustive it searches for their identities, while in the Raeng/Mascot approach the suitability of Mascot to find candidate sequences for α and β under the given conditions may by limiting.

### 3.7. New PPIs detected by pLink 2

The new cross-links identified by pLink 2 revealed 6 PPIs not identified in the Raeng/Mascot approach. One cross-linked peptide pair points to the interaction by which enzyme I (PT1_BACSU) from the sugar PTS system transfers a phosphoryl group from phosphoenolpyruvate (PEP) to the phosphoryl carrier protein HPr (PTHP_BACSU) [24]. Another cross-link reveals an interaction between the RNA chaperone CspB (CSBP_BACSU) and ribosomes, in particular with the 50S ribosomal protein L7/L12 (RL7_BACSU). Previously we found already a cross-link between CspB and the 30S ribosomal protein S2 [9]. This shows that CspB acts in close proximity of ribosomes, corroborating other observations, although a direct interaction with ribosomes could not be demonstrated before [25]. Four cross-links point to novel PPIs (Table 4). In supplementary Fig. 2 the corresponding mass spectra are depicted. One PPI not reported up to now as far as we know is revealed by a cross-linked peptide pair from the transition state regulatory protein AbrB (ABRB_BACSU) and translation elongation factor Tu (EFTU_BACSU). AbrB is known for its DNA binding activity of a large number of sites [26,27]. It can be phosphorylated by several protein kinases and it interacts with the anti-repressor AbbA [28–30]. The physiological significance of the interaction of AbrB with elongation factor Tu is not clear. AMPA_BACSU is a cytosolic aminopeptidase with broad specificity [31]. The interaction with the 50S ribosomal protein L17 (RL17_BACSU), which is located in close vicinity of the site where the nascent polypeptide emerges form the ribosome at the end of the tunnel [32] raises the question whether AmpA plays a role in either the removal of aminoterminal (formyl)-methionine during protein synthesis along with the canonical methionine aminopeptidases MAP and YflG [33,34] or in another post-translational modification. The only inter-protein cross-linked peptide pair of which one of the proteins (YopJ) is not a member of the 673 independently identified proteins, revealed an interaction with the α-subunit of RNA polymerase. The function of YopJ, a SPbeta prophage-derived protein, is not known. An intriguing interaction is found between the essential protein YlaN (YLAN_BACSU) [35] and the ferric uptake repressor Fur (FUR_BACSU) [36]. The function of YlaN has long been an enigma, but recent data point to a role in FeS cluster biogenesis [37]. The interaction of Ylan with Fur as shown here may provide a clue to understand how the effect of Ylan on FeS cluster biogenesis is brought about.

**Table 4.**
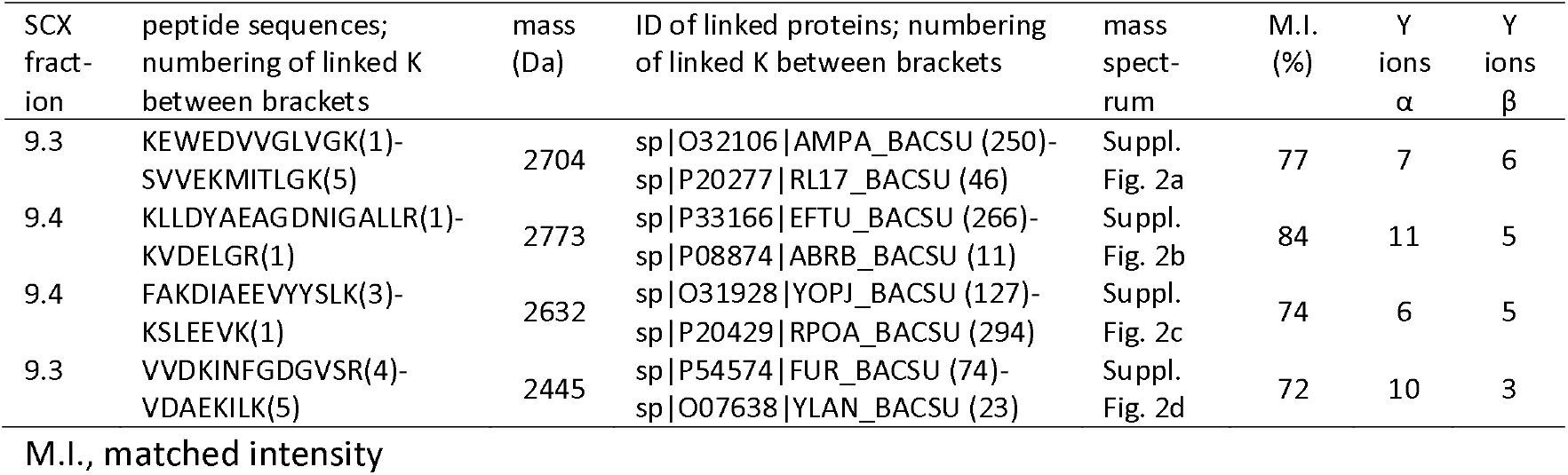
Cross-linked peptide pairs identified by pLink 2 revealing new protein-protein interactions

### 3.8. Contributions of the different assignment criteria to obtain a low FDR for inter-protein peptide pairs

In Table 5the contributions are listed of the different criteria used to discriminate between true and false positives. Details are depicted in Supplementary Tables S7 and S8. In the majority of cases the acceptance of assignment of a decoy sequence can only be prevented based on a single criterion, most decoy candidates passing the thresholds for the other criteria. The most discriminating criterion is the number of unambiguously assigned y ions. Unambiguousness in this respect is crucial, since it can significantly lower the FDR as shown in Table 5. The dependence of the elution time window during SCX chromatography on mass and charge of cross-linked peptide pairs is also a powerful criterion to identify a significant fraction of false positive inter-protein peptide pairs. SCX fractionation is often used for cleavable and non-cleavable cross-linkers using 1D [34–36] or 2D approaches [41]. The usually large population of intra-protein cross-linked peptide pairs can be used to prepare a set of references for SCX elution time windows. Here and previously [11,12] we have shown that the chance that intra-protein cross-link pairs are found by accident by searching an entire species specific sequence database is so small that false positives will be rare in this category of cross-links.

**Table 5.** Contribution of different criteria used to obtain a low FDR for inter-protein cross-linked peptide pairs

| Criterion | Decisive contribution to rejection of hits |  |  |  |
| --- | --- | --- | --- | --- |
|  | decoy hits |  | target hits |  |
|  | n | percent | n | percent |
| Matched intensity | 1 | 1.7 | 5 | 6.9 |
| Required number of y ions | 16 | 27.1 | 18 | 25.0 |
| Required number of b ions | 2 | 3.4 | 3 | 4.2 |
| Unambiguous assignment to y or b ions | 6 | 10.2 | 8 | 11.1 |
| Expected SCX elution time window | 5 | 8.5 | 7 | 9.7 |
| Combination of two or more of the above criteria | 22 | 37.3 | 26 | 36.1 |
| Better scoring candidate with the same mass and elution time in LCMSMS | 3 | 5.1 | 3 | 4.2 |
| Correction for formylation or carbamidomethylation | 3 | 5.1 | 2 | 2.8 |
| Assigned decoy sequence | 1 | 1.7 |  |  |
| Total | 59 |  | 72 |  |

### 3.9. A search at 0.1% FDR also requires additional filtering to obtain a low FDR for inter-protein peptide pairs

Finally the dataset was searched at 0.1% FDR by pLink 2 against the entire *B. subtilis* database. The data are depicted in Table 2. Slightly less candidates were nominated for both inter-protein peptide pairs, inter-protein peptide pairs and homo-dimer peptide pairs than in the search at 5% FDR. However, compared with the search at 5% FDR, exactly the same number and same identity of inter-protein peptide pairs were assigned after application of the composite filter. Only a few less candidates with aberrant SCX elution times or with one or both peptides not belonging to the 673 independently proteins were rejected by applying the composite filter than in the 5% FDR search. The relatively large number of rejected spurious candidates in a search with an overall FDR as low as 0.1%, underscores the usefulness of the composite filter to obtain a low FDR for inter-protein peptide pairs during interrogation of a large sequence database.

## 4. Conclusions

Here we show that pLink 2 efficiently nominated cross-linked peptide pairs from complex protein extracts after in vivo treatment of exponentially growing cells with BAMG. pLink 2 is also extremely fast as compared with use of our in house developed program Raeng for nomination of BAMG-cross-linked peptides. However, at an overall FDR of 5% or 0.1%, additional filtering is required to obtain a low FDR for inter-protein cross-linked peptide pair identifications. This is due to the notion that false positives are practically all confined to inter-protein peptide pairs when a search space as large as an entire species specific database is interrogated at a given FDR, if equal criteria for assignment of intra-protein and inter-protein peptides are used [11]. This can be circumvented by applying more stringent criteria for inter-protein cross-linked peptide pairs than for intra-protein cross-links. Here we show that matched intensity, the number of y ions to be assigned for both α and β and the unambiguousness of y ions, and to a lesser extent, b ions are useful criteria to diminish the FDR for inter-protein cross-linked peptide pairs. For future use of pLink 2 as a search engine for BAMG-cross-linked peptides pairs it would be useful if these criteria could be implemented in the pLink 2 code for cross-link approval. Also the number of assigned b ions may be included in the filter criteria, but with a correction factor for the relatively low average abundance as compared with the abundance of y ions. The isolation method used for BAMG-cross-linked peptides, diagonal SCX chromatography, offers an additional criterion for filtering based on their mass and charge at the pH of the chromatographic mobile phase in relation to the elution time.

About 20% high scoring candidates identified previously with Raeng had escaped detection by pLink 2, and vice versa. While pLink 2 is fast and efficient, identifying approximately the same number of cross-linked-peptide pairs as compared with our previous approach, it would be worthwhile to understand the reasons for the slightly difference in output between these two approaches for the benefit of future research with BAMG and other cleavable cross-linkers.

## Acknowledgements

The authors thank PhD student Zhen-Lin Chen for adapting pLink 2 to enable the use of BAMG in the stepped HCD mode and for valuable comments during the preparation of the manuscript.

## Supplementary material

**Supplementary Table S1.**
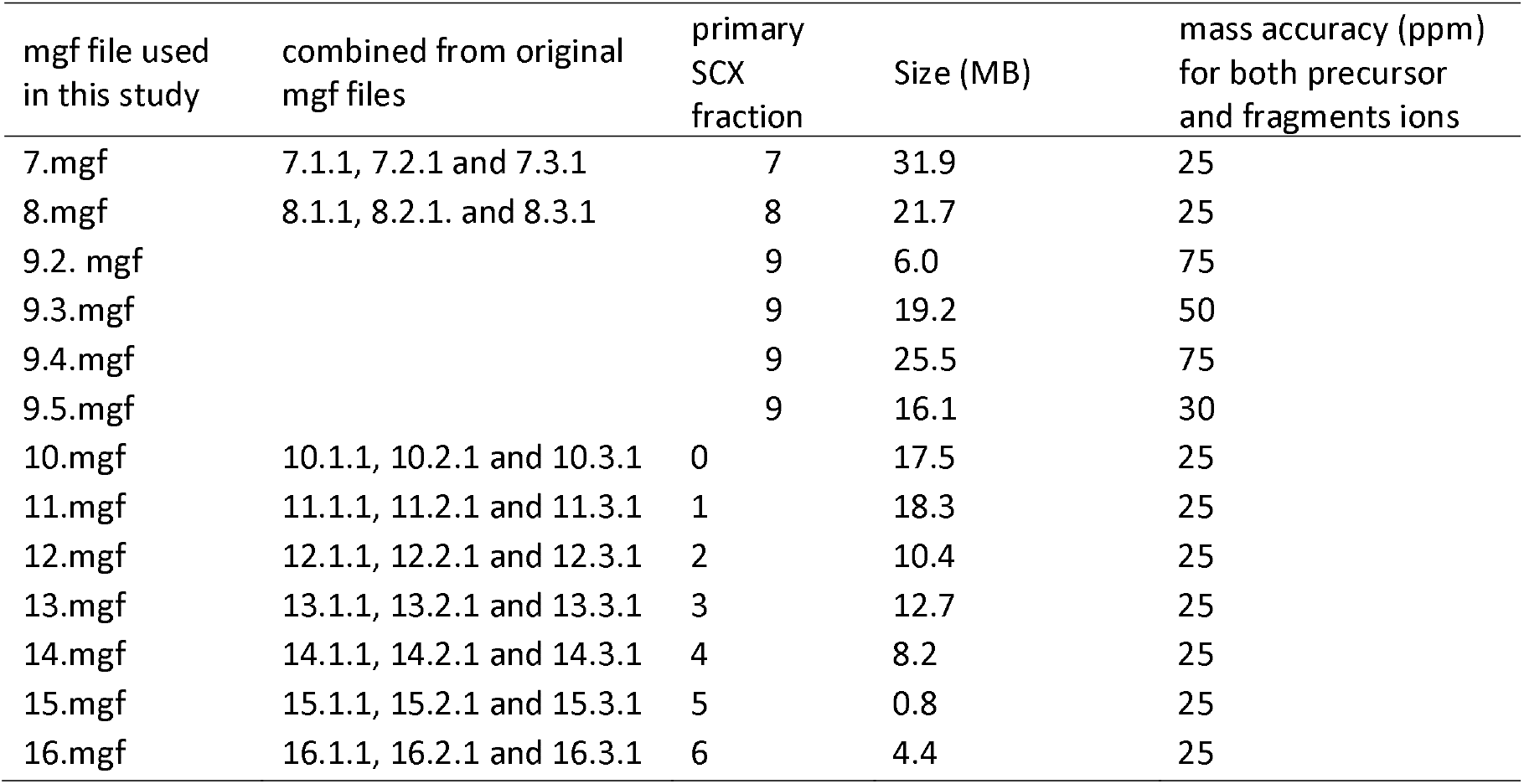
Mgf files used in this study

### Legends to supplementary figures

**Supplementary figure 1.**
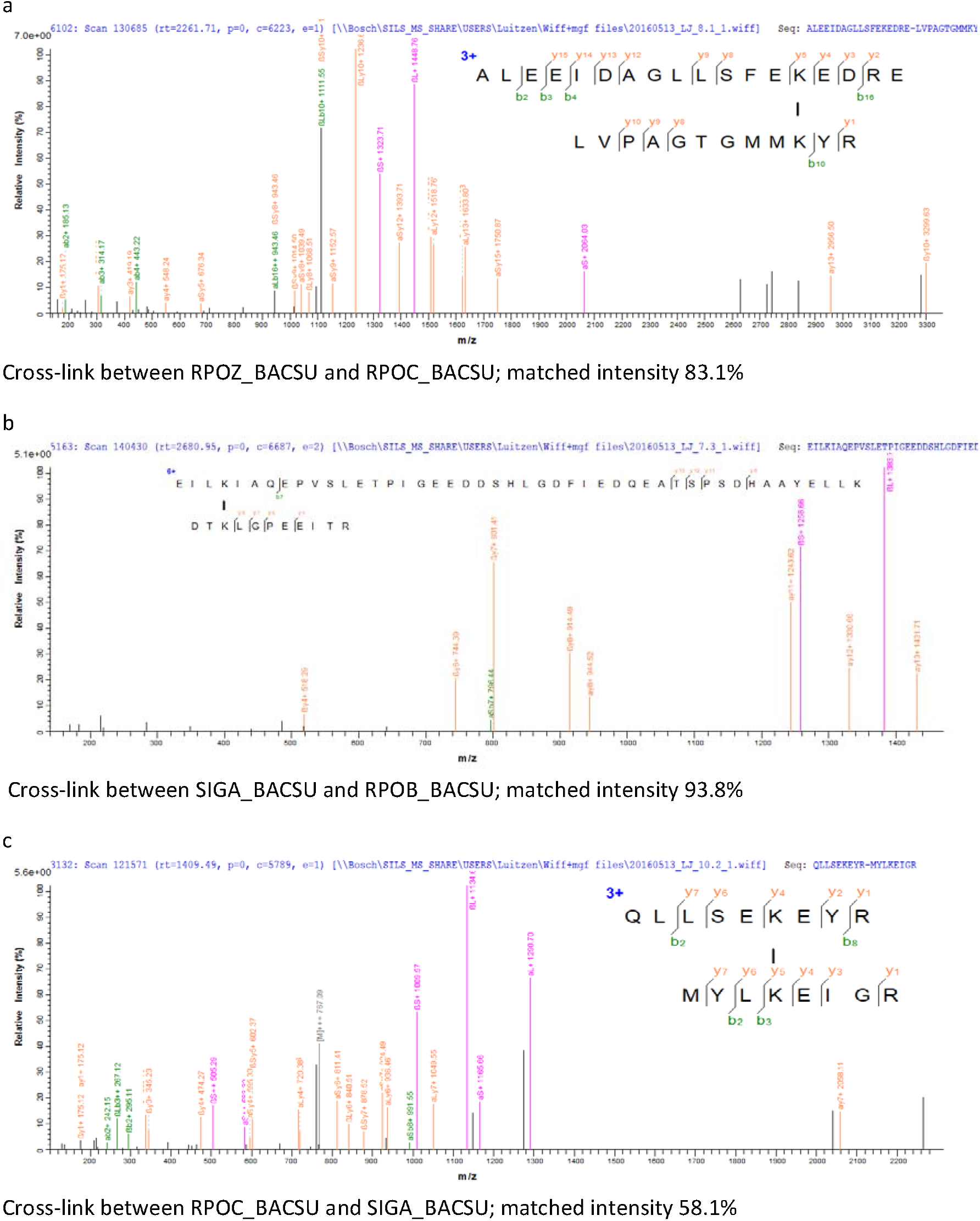
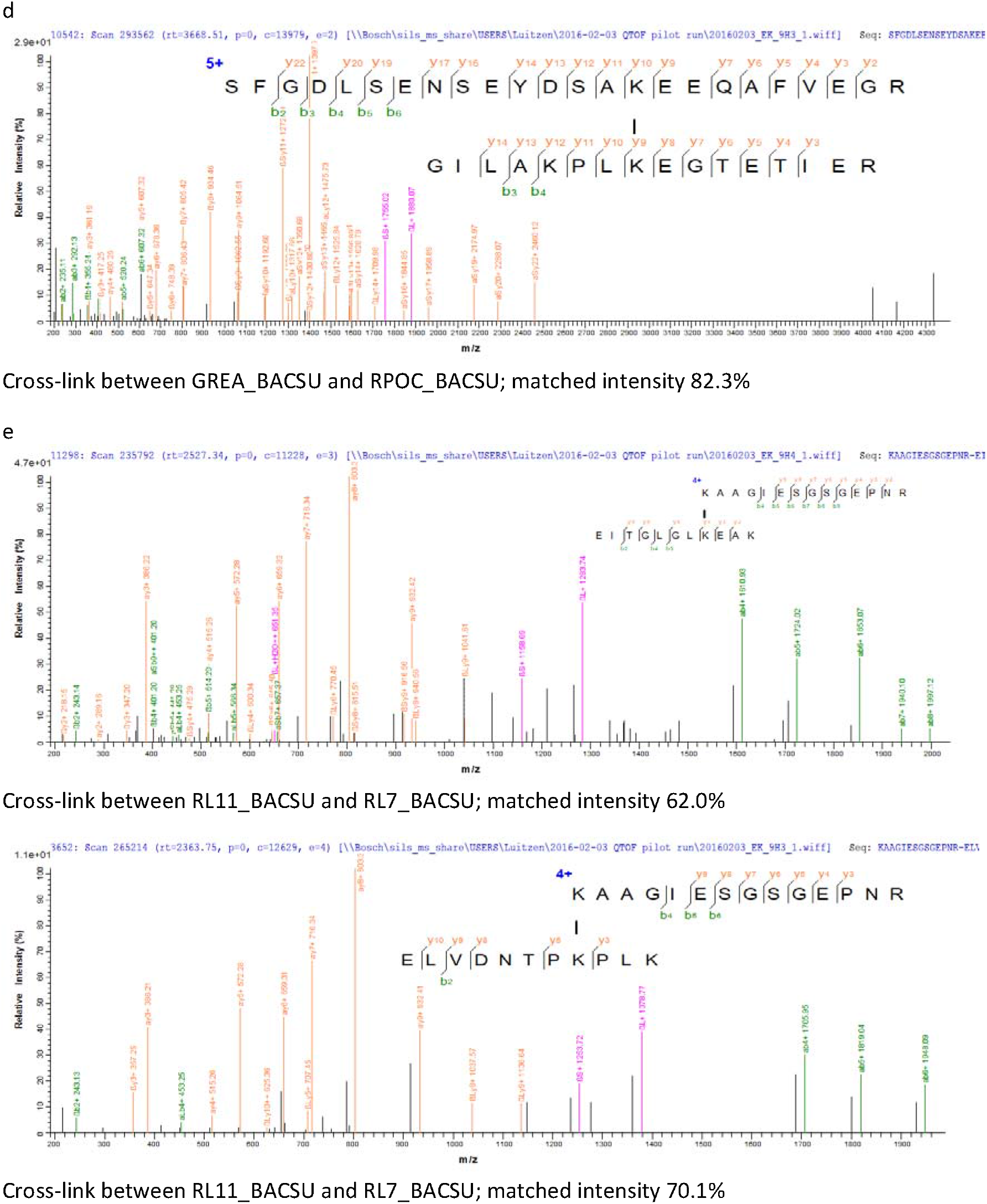
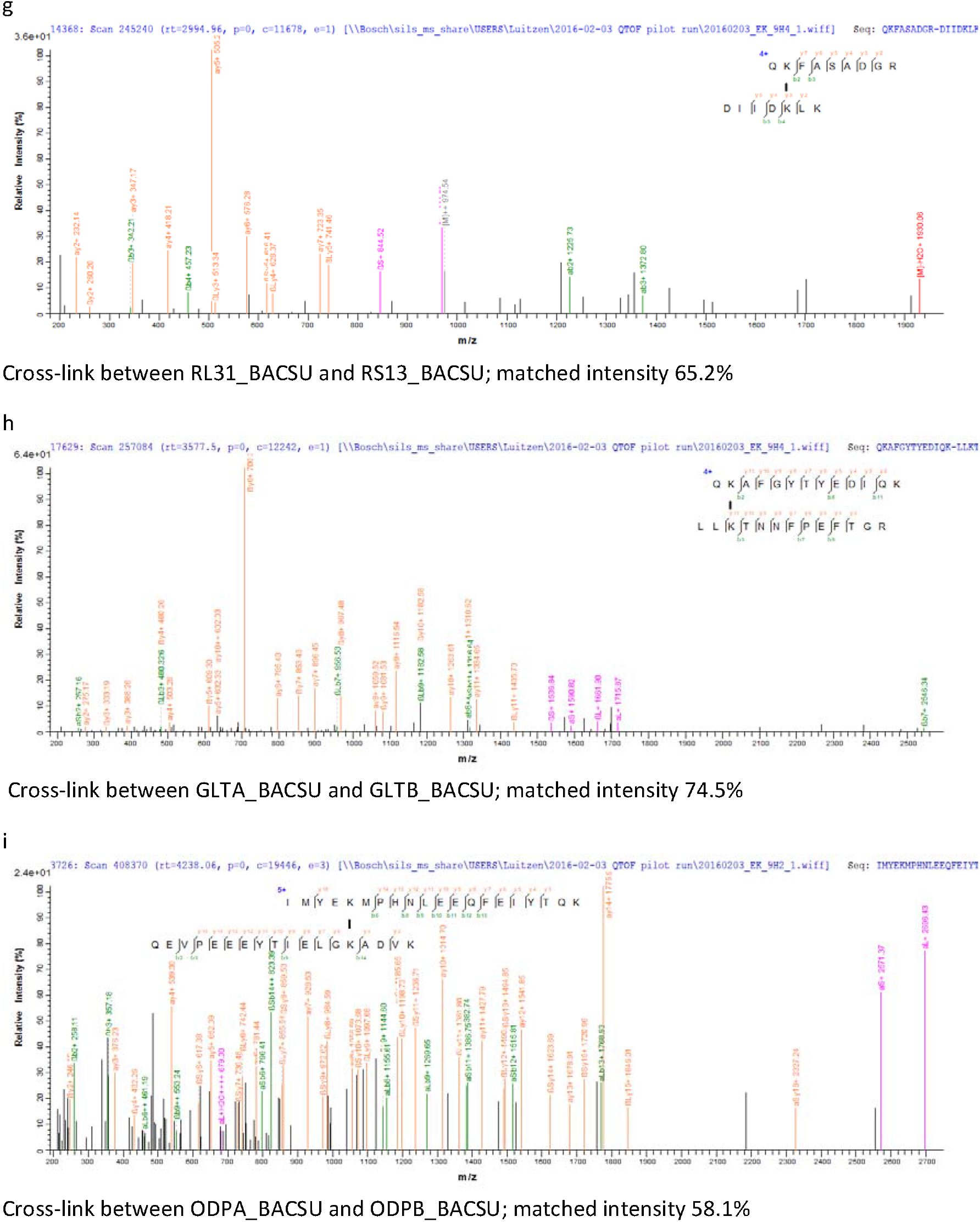
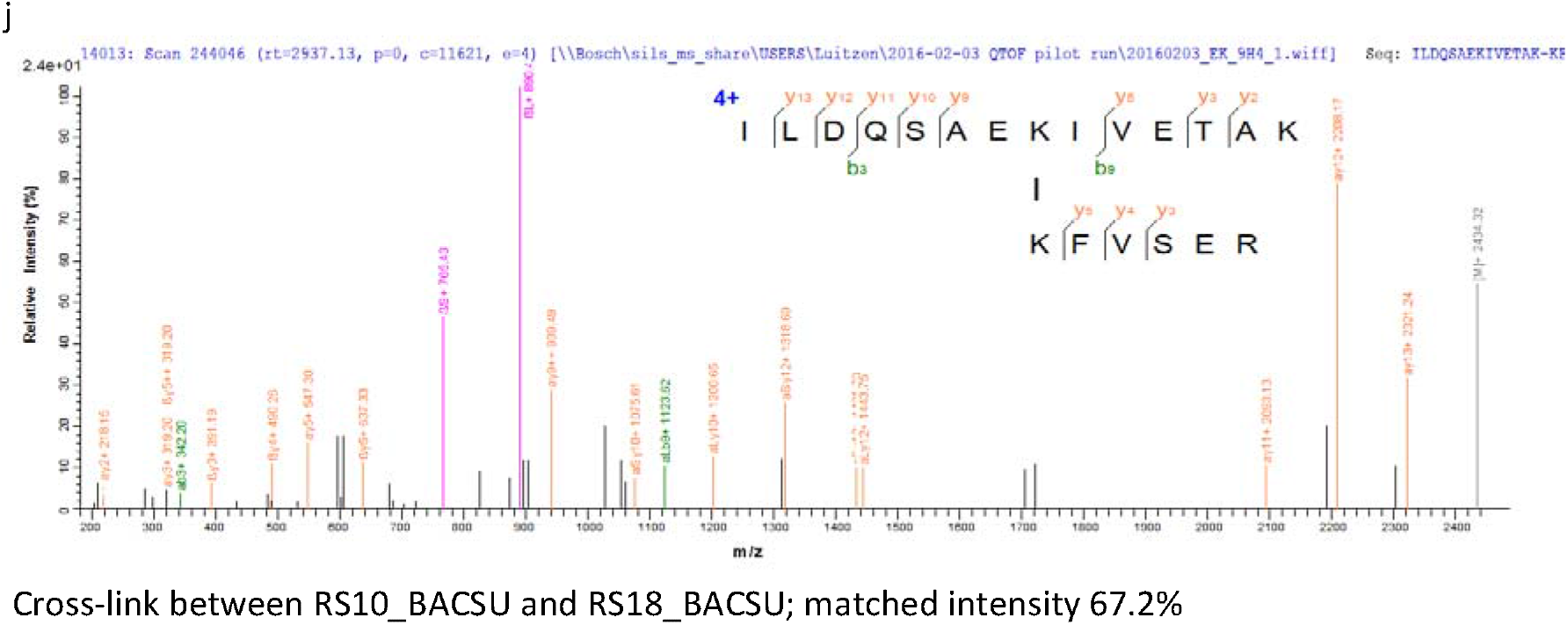
Mass spectra with high scores of inter-protein cross-linked pairs identified by Raeng/Mascot, but not pLink 2 showing interactions between a, RPOZ_BACSU and RPOC_BACSU; b, SIGA_BACSU and RPOB_BACSU; c, RPOB_BACSU and SIGA_BACSU; d, GREA_BACSU and RPOC_BACSU; e, RL11_BACSU and RL7_BACSU; f, RL11_BACSU and RL7_BACSU; g, RL31_BACSU and RS13_BACSU; h, GLTA_BACSU and GLTB_BACSU; i, ODPA_BACSU and ODPB_BACSU; j, RS10_BACSU and RS18_BACSU

**Supplementary figure 2.**
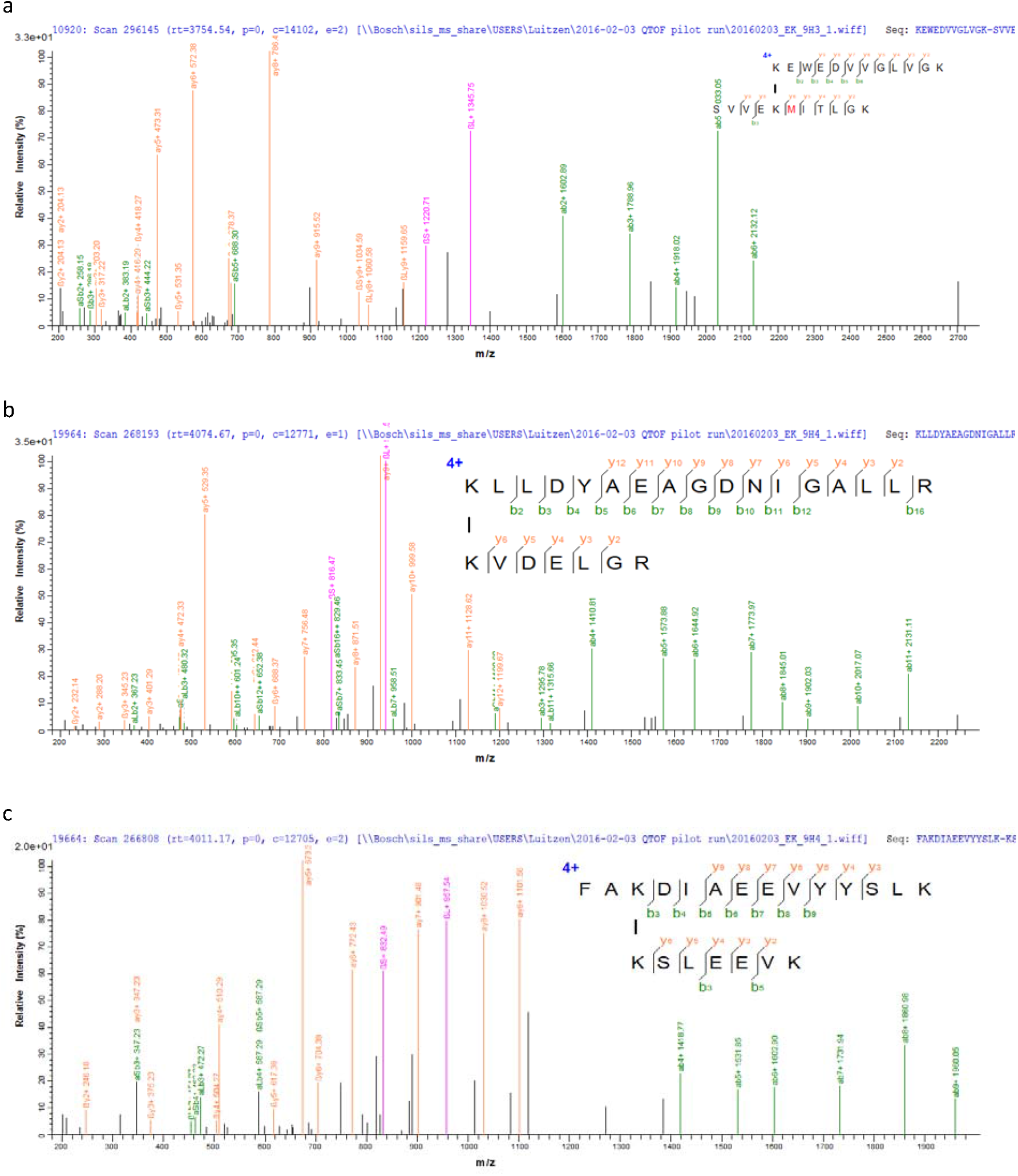
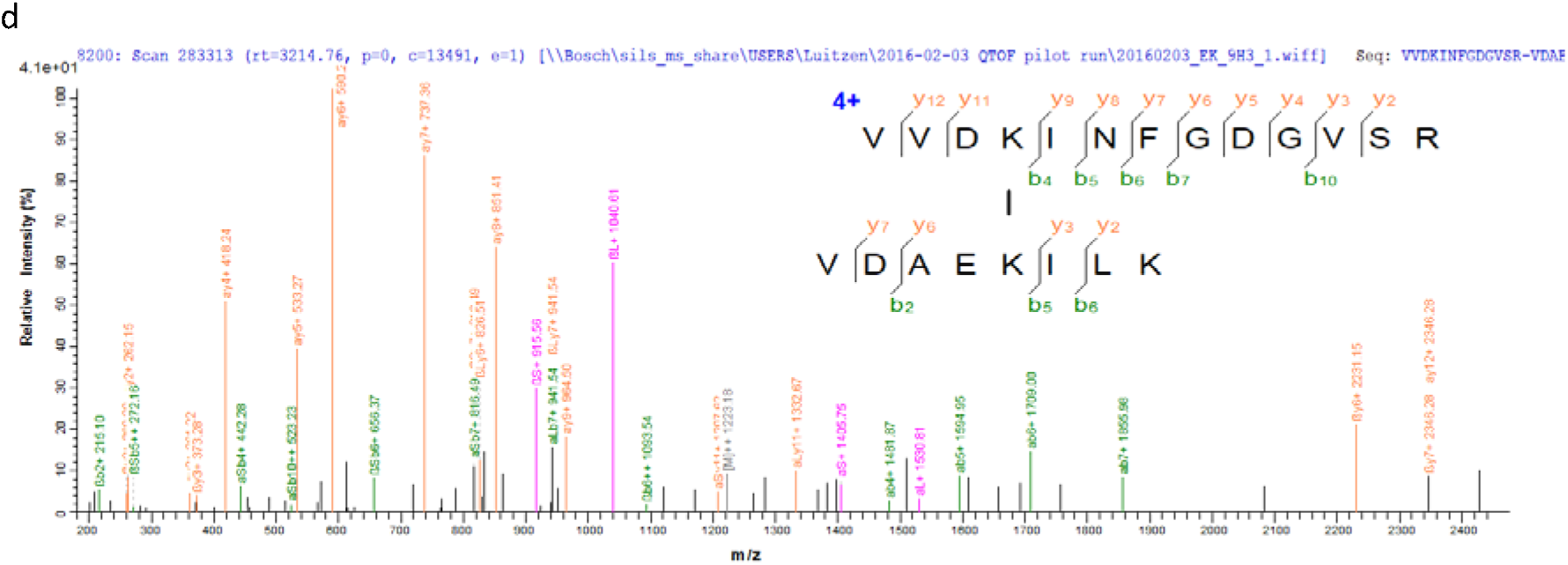
Mass spectra of cross-linked peptide pairs identified by pLink 2 revealing new protein-protein interactions a, AMPA_BACSU-RL7_BACSU; b, EFTU_BACSU-ABRB_BACSU; c, YOPJ_BACSU-RPOA_BACSU; d, FUR_BACSU-YLAN_BACSU

